# Human cytomegalovirus protein RL1 degrades the antiviral factor SLFN11 via recruitment of the CRL4 E3 ubiquitin ligase complex

**DOI:** 10.1101/2021.05.14.444170

**Authors:** Katie Nightingale, Ceri A. Fielding, Cassie Zerbe, Leah Hunter, Martin Potts, Alice Fletcher-Etherington, Luis Nobre, Eddie C.Y. Wang, Blair L. Strang, Jack Houghton, Robin Antrobus, Nicolas M. Suarez, Jenna Nichols, Andrew J. Davison, Richard J. Stanton, Michael P. Weekes

**Author notes:** Corresponding author, **Correspondence:** Michael P. Weekes. **Author Contributions:** KN and MPW designed the research. KN, CAF, CZ, LH, MP, AFE, LN, ECYW, RA, NMS and JN performed the research. KN, JH, BLS, AJD and MPW analysed the data. AJD, RJS and MPW supervised the research. KN and MPW wrote the manuscript. All authors edited the manuscript. **Competing Interest Statement:** No competing interests.

## Abstract

Human cytomegalovirus (HCMV) is an important human pathogen and a paradigm of viral immune evasion, targeting intrinsic, innate and adaptive immunity. We have employed two novel, orthogonal multiplexed tandem mass tag-based proteomic screens to identify host proteins downregulated by viral factors expressed during the latest phases of viral infection. This approach revealed that the HIV-1 restriction factor Schlafen-11 (SLFN11) was degraded by the poorly characterised, late-expressed HCMV protein RL1, via recruitment of the Cullin4-RING E3 Ubiquitin Ligase (CRL4) complex. SLFN11 potently restricted HCMV infection, inhibiting the formation and spread of viral plaques. Overall, we show that a restriction factor previously thought only to inhibit RNA viruses additionally restricts HCMV. We define the mechanism of viral antagonism and also describe an important resource for revealing additional molecules of importance in antiviral innate immunity and viral immune evasion.

**Significance Statement:** Previous proteomic analyses of host factors targeted for downregulation by HCMV have focused on early or intermediate stages of infection. Using multiplexed proteomics, we have systematically identified viral factors that target each host protein downregulated during the latest stage of infection, after the onset of viral DNA replication. Schlafen-11 (SLFN11), an interferon-stimulated gene and restriction factor for retroviruses and certain RNA viruses, potently restricted HCMV infection. Our discovery that the late-expressed HCMV protein RL1 targets SLFN11 for proteasomal degradation provides the first evidence for a viral antagonist of this critical cellular protein. We therefore redefine SLFN11 as an important factor that targets DNA viruses as well as RNA viruses, offering novel therapeutic potential via molecules that inhibit RL1-mediated SLFN11 degradation.

## Introduction

Human cytomegalovirus (HCMV) is a ubiquitous pathogen that establishes a lifelong latent infection in the majority of the world’s population (1). Reactivation from latency in immunocompromised individuals, such as transplant recipients and AIDS patients, can result in significant morbidity and mortality (2). HCMV is also the leading cause of infectious congenital birth defects, including deafness and intellectual disability, affecting ~1/100 pregnancies (1). However, only a few antiviral drugs are approved for the treatment of HCMV, all of which are associated with significant toxicity, and there is currently no licensed vaccine (3).

Susceptibility to viral infection and disease is determined in part by antiviral restriction factors (ARFs) and the viral antagonists that have evolved to degrade them (4). Small molecules that inhibit ARF-antagonist interactions may restore endogenous restriction and offer novel therapeutic potential (5). Identification of novel ARFs and characterisation of their interactions with HCMV antagonists is therefore clinically important.

HCMV possesses the largest human herpesvirus genome, encoding 170 canonical open reading frames (ORFs). A modest number of non-canonical ORFs may encode additional functional proteins (6–9). During productive HCMV infection, viral gene expression occurs in cascades during a ~96 h infection cycle that is conventionally divided into immediate-early, early and late phases. Early genes encode functions necessary for initiating viral DNA replication. In the late phase, early-late genes are initially transcribed at low levels and are then upregulated after the onset of viral DNA replication, whereas true-late genes are expressed exclusively after DNA replication commences and include proteins required for HCMV virion assembly. We previously characterised five temporal classes of viral protein expression, offering finer definition of protein expression profiles (10).

As over 900 proteins are downregulated more than three-fold during the course of HCMV infection, predicting molecules likely to perform novel immune functions is challenging without additional data (7, 10, 11). Our previous analysis of the subset of proteins targeted for degradation by 24 or 48 h led directly to the identification of Helicase-Like Transcription Factor (HLTF) as a novel ARF, and HCMV UL36 as a key inhibitor of necroptosis, by degrading Mixed Lineage Kinase-domain-Like protein (MLKL) (7, 10). However, no studies have systematically examined which host factors are targeted by viral proteins during the latest phase of infection. This question is important as some host factors may play important roles in restricting the final stages of viral replication. Furthermore, despite our prior characterisation of a comprehensive HCMV interactome (9), the abundance of certain host proteins whose expression is downregulated during infection can be sufficiently low to impede identification of their viral antagonists.

We have used two complementary proteomic approaches to address these questions. The first identified cellular proteins specifically targeted by HCMV factors expressed after viral DNA replication, by comparing host protein expression over time in the presence or absence of the viral DNA polymerase inhibitor phosphonoformic acid (PFA). The second employed an enhanced panel of HCMV mutants each deleted in contiguous gene blocks dispensable for virus replication *in vitro*, most of which we have described previously (12).

The intersection between these approaches showed that one particular protein, Schlafen family member 11 (SLFN11), is both downregulated during the late phase of HCMV infection and is targeted by the RL1-6 block of viral genes. SLFN11 potently restricted HCMV infection and therefore represents a novel HCMV ARF. Among the factors encoded by the RL1-6 region, RL1 was required for SLFN11 downregulation, via recruitment of the Cullin4-RING E3 Ubiquitin Ligase (CRL4) complex. Overall, our data identifies a novel HCMV ARF and a novel mechanism of viral antagonism, and describes an important resource that will reveal additional molecules of importance in antiviral innate immunity and viral immune evasion.

## Results

### Host proteins downregulated by late-expressed HCMV factors

To globally quantify cellular proteins whose expression is increased or decreased by late-expressed HCMV factors, we applied PFA to HCMV-infected primary human fetal foreskin fibroblasts (HFFFs) at the time of infection and harvested samples for analysis at 24h intervals (**Figure 1A**). Expression of early viral genes is largely unaffected by PFA, whereas early-late genes are partially inhibited and late genes are completely inhibited (13). Ten-plex tandem mass tag (TMT) technology and MS/MS/MS mass spectrometry of wholecell lysates enabled precise protein quantification (**Figure 1A**).

**Figure 1.**
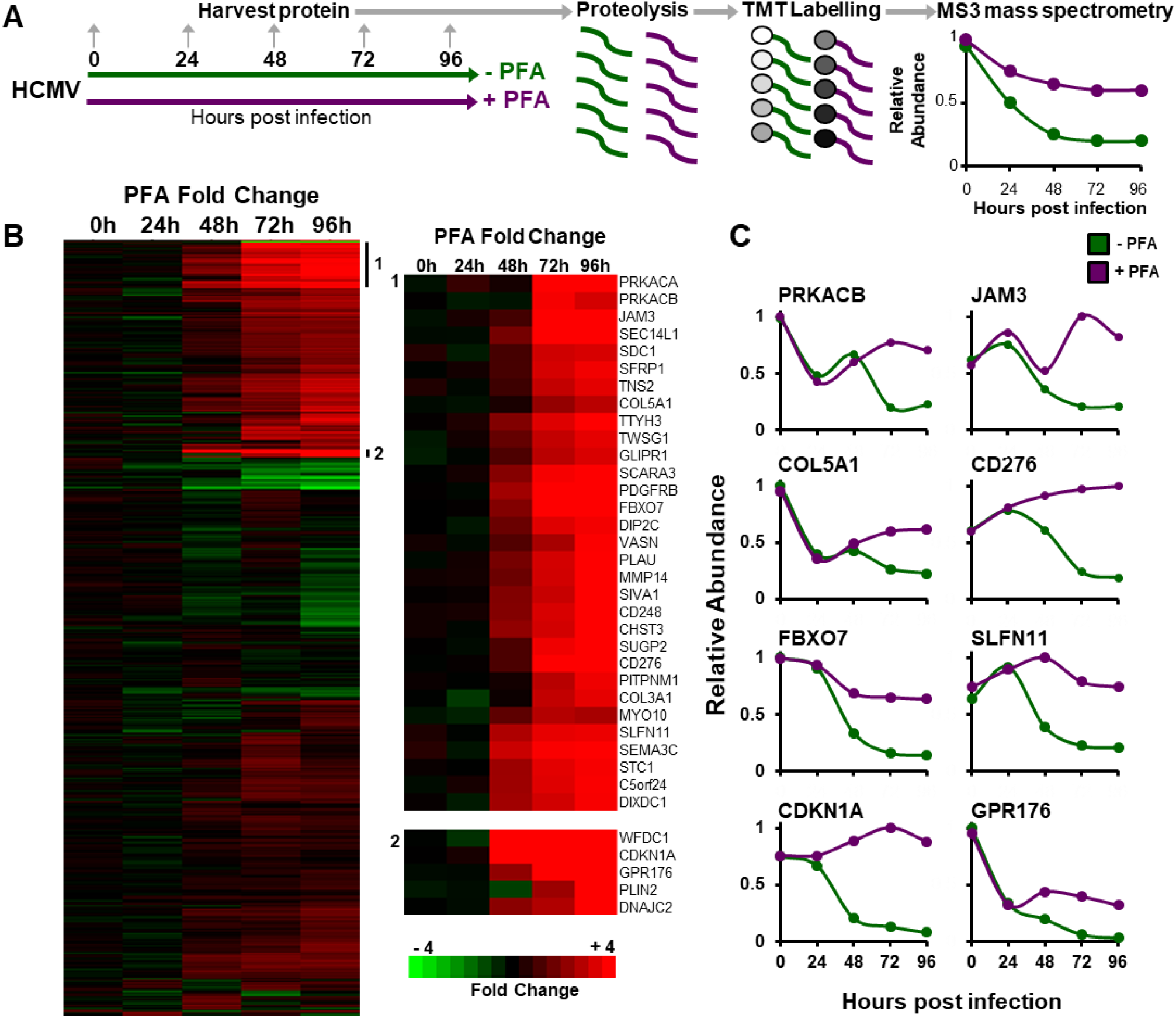
Host proteins targeted for downregulation by HCMV late during infection, identified using the viral DNA synthesis inhibitor PFA. (A) Schematic of the experimental workflow. HFFFs were infected with HCMV at multiplicity of infection (MOI) 10, and cells were harvested at the indicated times. (B) Hierarchical cluster analysis of 527 proteins downregulated ≥3-fold by 96 hpi. For each protein, the ratios of protein expression in the presence or absence of PFA are shown. To be considered a ‘hit’ in the screen, proteins were additionally required to be rescued >2-fold by PFA. Enlargements to the right of the panel show examples of subclusters. (C) Examples of temporal profiles of proteins rescued from downregulation by PFA.

We quantified 8059 human and 149 viral proteins, and observed good correspondence between proteins modulated during HCMV infection in the absence of PFA and protein expression in our previously published proteomic datasets (10) (**Figure S1**). Overall, by 96 hours post infection (hpi), 157 human proteins were downregulated ≥3-fold in the absence of PFA and ‘rescued’ >2-fold in the presence of PFA (**Figure 1B, Dataset S1A**). Application of DAVID software (14) indicated that these included groups of plasma membrane proteins, proteins with immunoglobulin or cadherin domains, and proteins with functions in viral infection (**Figure S2A, Dataset S1B**). Examples included multiple collagens, ephrins, syndecans and adhesion molecules such as junctional adhesion molecule-3 (JAM3), in addition to T-cell co-stimulator CD276 and DNA replication inhibitor and HIV-1 restriction factor Schlafen-11 (SLFN11) (15, 16) (**Figure 1C**). Additionally, 87 human proteins were both upregulated ≥3-fold by 96 hpi yet downregulated >2-fold in the presence of PFA (**Figures S2B-C, Dataset S1C**), indicating that late-expressed viral proteins can exhibit additional functions in host regulation.

**Figure S1.**
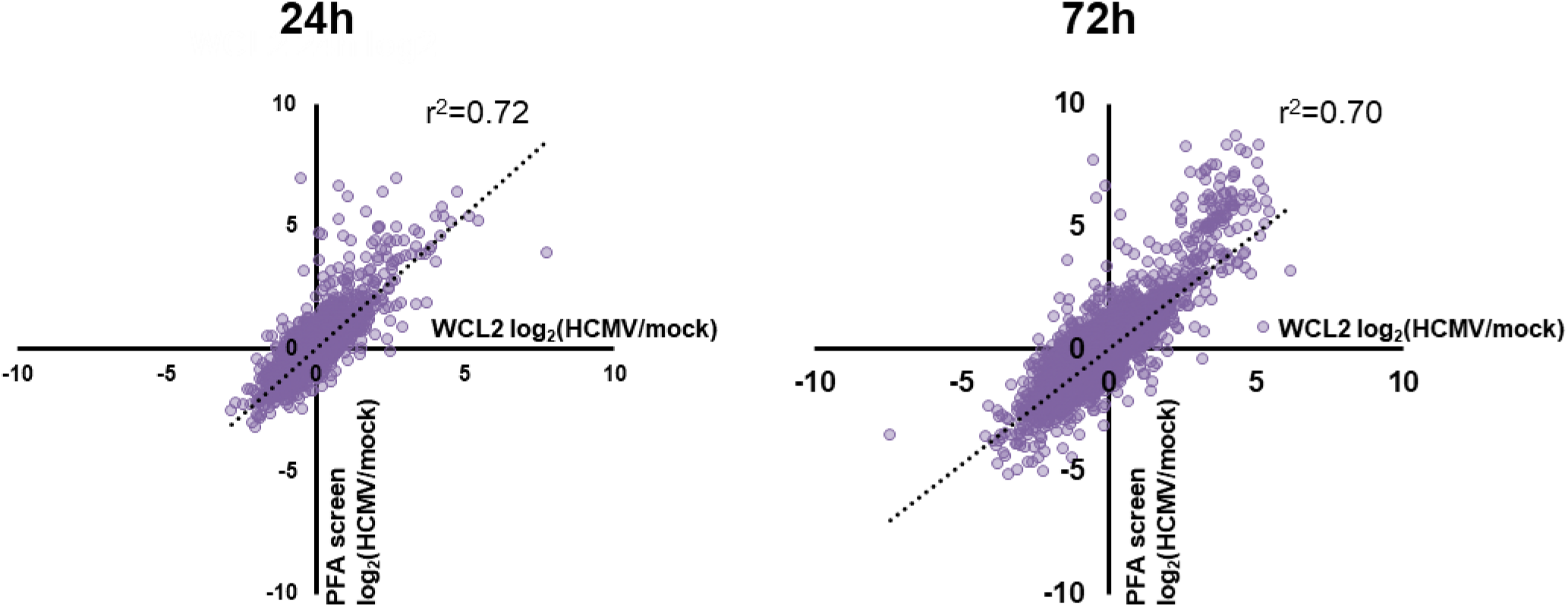
Comparison between data from our previously published temporal analysis of HCMV infection (experiment ‘WCL2’), and the arm of the present experiment (‘PFA screen’) without PFA treatment. Cells were infected with HCMV or mock-infected for 24 or 72 h. HCMV:mock ratios for each protein were highly correlated between experiments.

**Figure S2.**
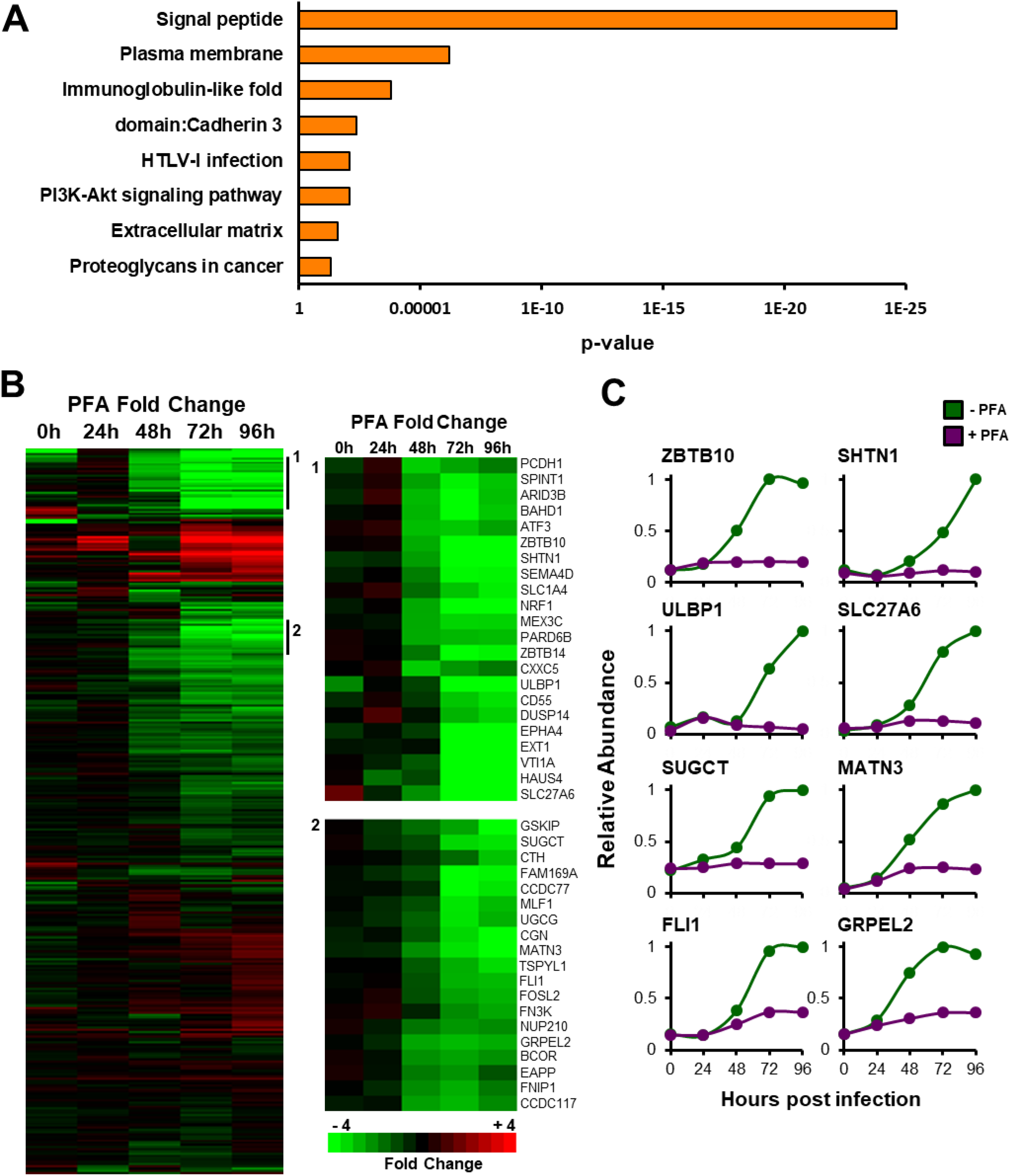
(A) DAVID analysis of pathway enrichment among ‘hits’ in the PFA screen (proteins downregulated ≥3-fold by HCMV over the course of infection and rescued >2-fold by the addition of PFA). Benjamini-Hochberg adjusted p-values are shown for each pathway. (B) Hierarchical cluster analysis of 332 human proteins upregulated ≥3-fold by HCMV over the course of infection. For each protein, the ratios of protein expression in the presence or absence of PFA are shown. Enlargements to the right of the panel show examples of subclusters. (C) Examples of temporal profiles of proteins whose expression was downregulated in the presence of PFA.

### RL1 is necessary and sufficient for SLFN11 downregulation

Identification of which HCMV protein(s) target a given cellular factor can be challenging due to the substantial coding capacity of HCMV. To identify viral proteins targeting host factors late during HCMV infection, we extended our previous approach that analysed infection at 72 h with a panel of recombinant viruses, each deleted for one or other of a series of blocks of genes non-essential for replication *in vitro* (12) (**Figure S3A**). In this analysis, all viruses were examined in at least biological duplicate, and for the first time ΔRL1-6 HCMV was included since the functions of HCMV factors encoded within this gene block (RL1, RL5A, RL6 proteins and the RNA2.7 long non-coding RNA) are poorly characterised. For each human protein, a z-score and fold change (FC) compared to wild-type (wt) infection was calculated (see Materials and Methods). Sensitive criteria with a final z-score of >4 and FC >1.5 assigned 254 modulated cellular proteins to viral blocks (**Figure 2A**), and stringent criteria (z-score>6, FC>2) assigned 109 proteins to viral blocks (**Figure S3B, Dataset S2**). Data from this and the PFA screens are shown in **Dataset S3**, where the worksheet ‘‘Plotter’’ is interactive, enabling generation of graphs of expression of any of the human and viral proteins quantified.

**Figure S3.**
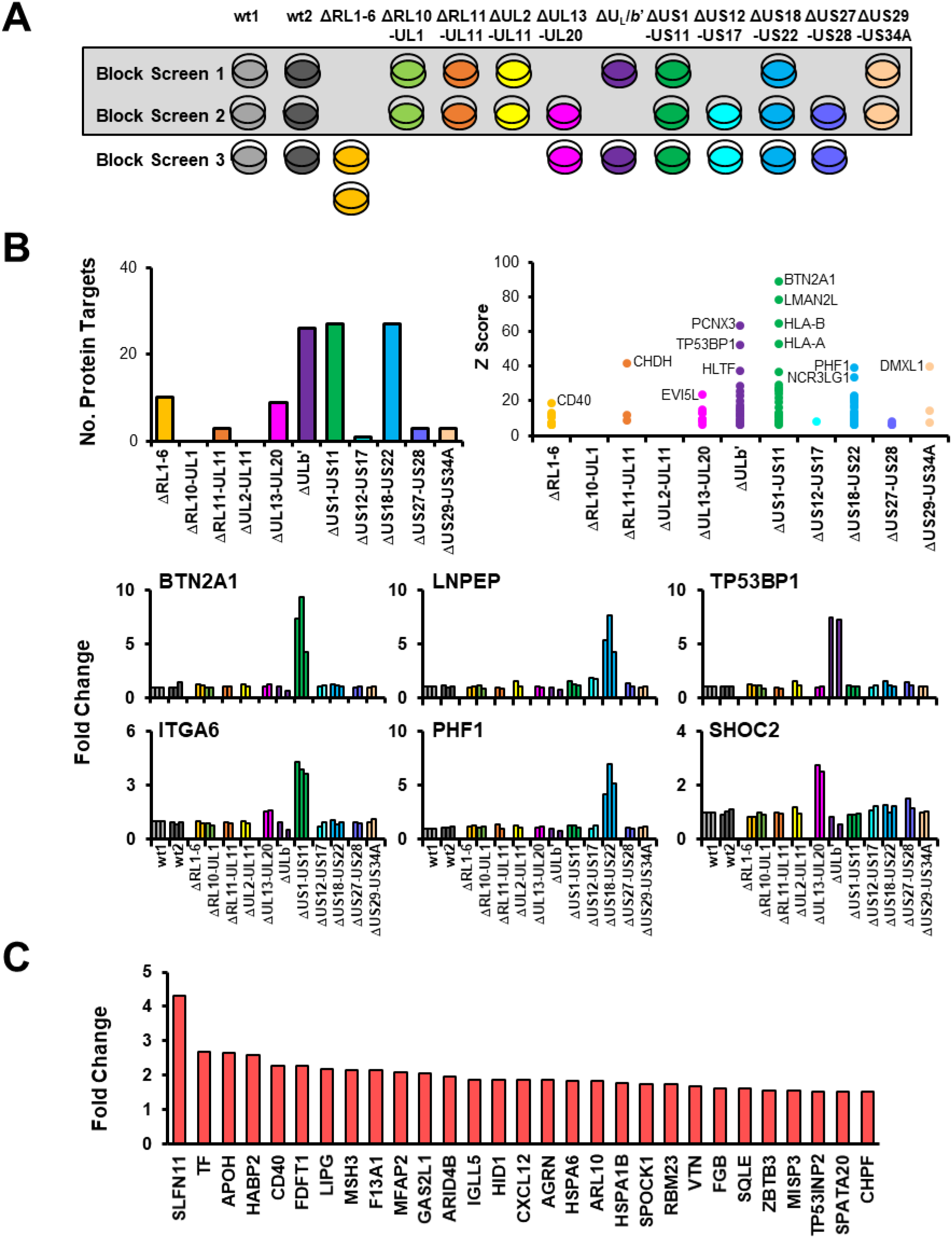
(A) Schematic of the gene block deletion experiment. Experiments 1 and 2 were reported previously (7), and experiment 3 was performed for the present study, for the first time including the ΔRL1-RL6 gene block deletion mutant, in biological duplicate. (B) (top left panel) Numbers of human proteins targeted by each block using stringent scoring (z-score >6 and fold change >2). For each block, the z-scores of all proteins that passed scoring criteria are shown (top right panel). Further details of the results using stringent scoring are shown in **Figure 2A** and in the Materials and Methods. (bottom panels) Further examples of results (see also **Figure 2C**). (C) Bar chart illustrating proteins ‘rescued’ upon deletion of the RL1-RL6 gene block. Fold change values shown were calculated by (average signal : noise (RL1-RL6 gene block deletion mutant) / signal : noise (parental HCMV)). Each protein with a fold change of >2 is illustrated.

**Figure 2.**
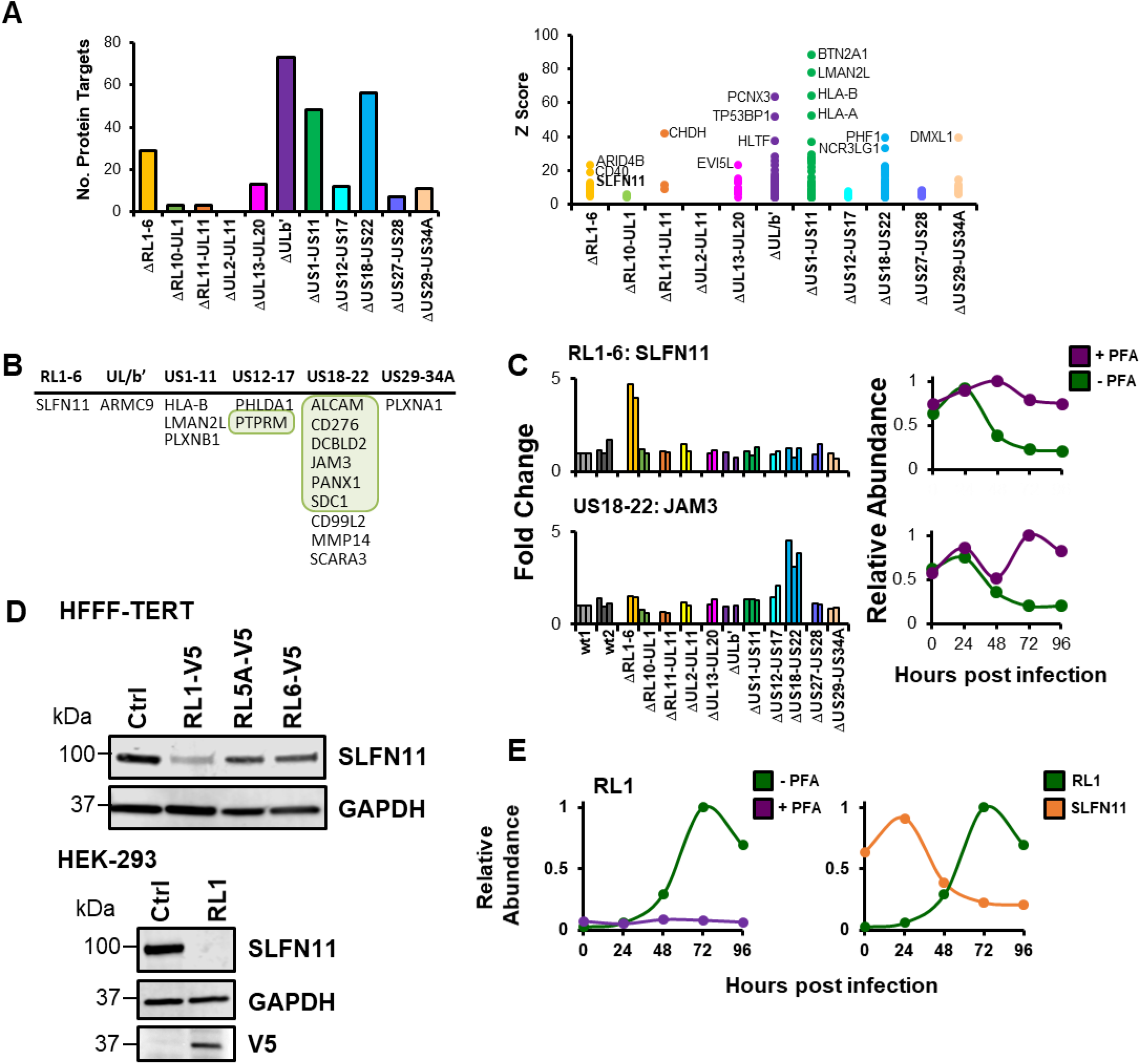
HCMV RL1 is necessary and sufficient for downregulation of SLFN11. (A) (left panel) Numbers of human proteins targeted by each gene block using sensitive scoring (z-score >4 and FC >1.5). For each block, the z-scores of all proteins that passed scoring criteria are shown (right panel). All viruses were examined in duplicate or triplicate across three separate experiments, the first two of which we have published previously (7) (**Figure S3A**). Infection was at MOI 10 for 72 h. Further details are given in Materials and Methods. (B) Table of 17 proteins that were downregulated >3-fold during HCMV infection, rescued >2-fold by PFA (**Figure 1B**), and passed sensitive scoring criteria to identify the targeting gene block. (C) Examples of data for proteins listed in (B). (D) Immunoblot confirming that RL1 alone is sufficient for downregulation of SLFN11 in stably transduced HFFF-TERTs (top panel) and transiently transfected HEK-293s (bottom panel). As we reported previously (9), expression of RL5A and RL6 was not detected by immunoblot, whereas both were detected by mass spectrometry (**Figure 3A, Dataset S4**). (E) Expression of RL1 is inhibited by PFA (left panel). The temporal profile of RL1 expression correlates inversely with expression of SLFN11 (right panel). Data for each protein is shown from the ‘PFA screen’ proteomic experiment (**Figure 1A**).

To identify host factors targeted for downregulation by late-expressed HCMV proteins, data from the PFA and gene-block screens were combined. Using sensitive criteria, 17 host proteins that were downregulated ≥3-fold by 96 hpi, ‘rescued’ >2-fold by PFA and targeted by one or other of the viral gene blocks examined (**Figure 2B**). These included proteins with previously described HCMV protein antagonists, for example known targets of the US18-US22 block including ALCAM, CD276 and JAM3, and PTPRM, which is a target of the US12-US17 block (**Figures 2B-C**) (17). The only assigned target of the RL1-RL6 block that met the threshold for rescue by PFA was SLFN11 (**Figure 2C**). Furthermore, of the proteins targeted this block, SLFN11 was the most substantially modulated (**Figure S3C**).

To determine which viral protein targets SLFN11 for downregulation, C-terminally V5-tagged RL1, RL5A and RL6 constructs were stably overexpressed in HFFFs immortalised with human telomerase (HFFF-TERTs). Overexpression of RL1-V5 alone was sufficient for downregulation of SLFN11 and this was recapitulated by transient transfection of HEK-293 cells with RL1-V5 (**Figure 2D**). Expression of RL1 was completely inhibited by the addition of PFA, and the profile of RL1 expression inversely correlated with the profile of SLFN11 (**Figure 2E**).

### RL1 degrades SLFN11 through recruitment of the Cullin4 E3 Ligase Complex

The HCMV RL1 and UL145 genes are related to each other and thus belong to the RL1 family (6). We and others have previously shown that the UL145 protein can employ CRL4 complex components CUL4A and DDB1 to degrade HLTF and STAT2 (7, 18). Using SILAC immunoprecipitation and coimmunoprecipitation, we identified a similar interaction between RL1 and DDB1 and CUL4A (**Figures 3A-B, Dataset S4**). A panel of alanine substitution mutations was tested to identify the region within RL1 required for interaction with DDB1 based on the DDB1 interaction motif previously identified within UL145 (19) (**Figure 3C**). As predicted, residues LL153-4, R157 and R159 were required for DDB1 binding, whereas residue T152 was dispensable. In contrast to residue N25 in UL145, which is indispensable for binding DDB1, the equivalent residue P149 in RL1 was not required. This may reflect the differences in the chemical properties of proline and asparagine residues, or the conservation within the DCAF family of asparagine at this position. Residues LL153-4, R157 and R159 are completely conserved across all publicly available HCMV RL1 sequences (263 different strains), and the corresponding residues in HCMV UL145 are also completely conserved (264 different strains) (unpublished data). Furthermore, the LLxxRxR motif is highly conserved (complete conservation in 7/8 RL1 orthologues and 8/8 UL145 orthologues, **Figure S4**).

**Figure 3.**
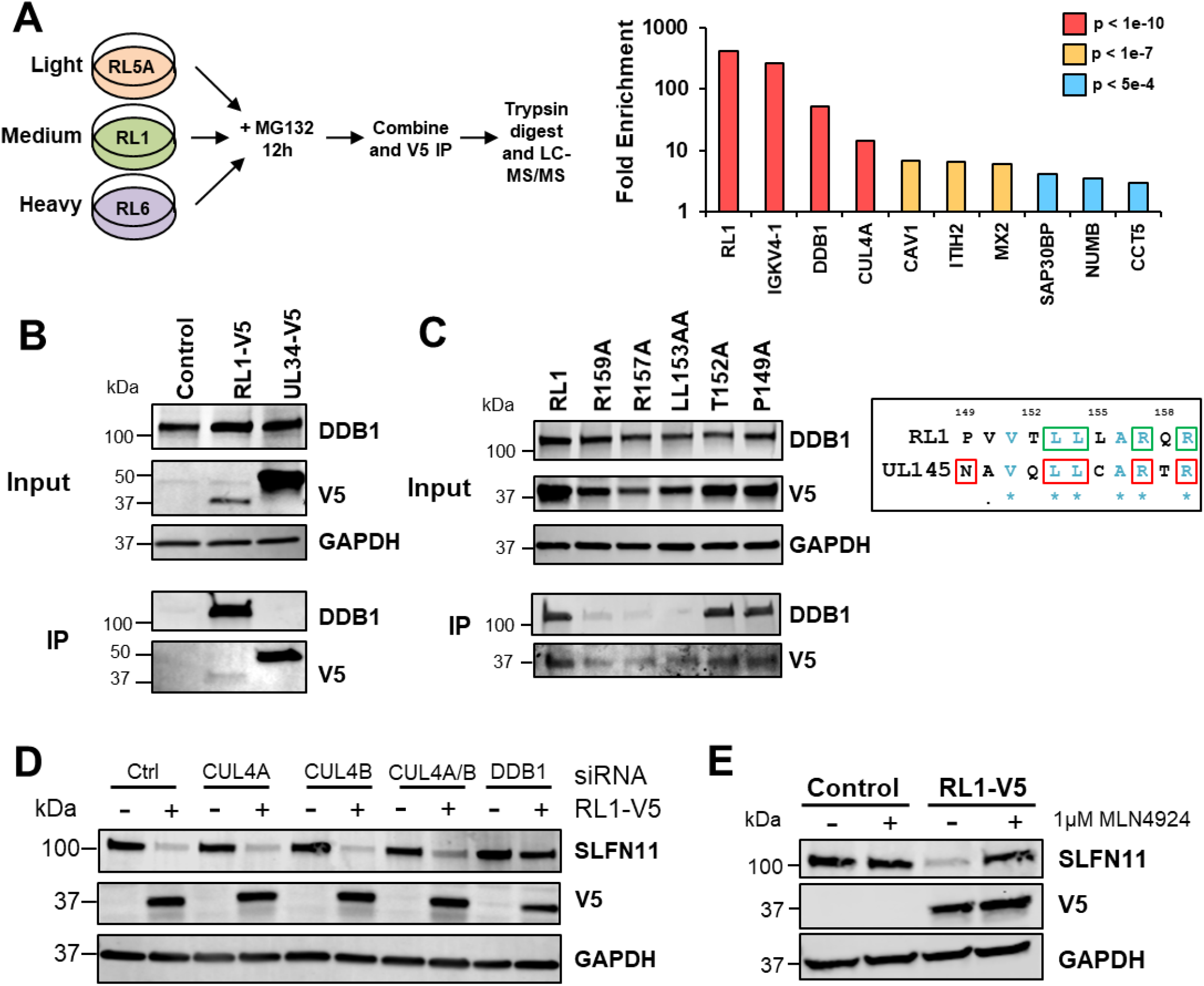
HCMV RL1 degrades SLFN11 via the CRL4 Complex. (A) (left panel) Schematic of SILAC immunoprecipitation. HFFF-TERTs stably transduced with C-terminally V5-tagged RL1 or RL5A or RL6 as controls were treated with 10 μM MG132 for 12 h prior to harvest. (right panel) Proteins enriched >3-fold in RL1-expressing cells compared with RL6-expressing cells are shown. p-values were estimated using significance A values, then corrected for multiple hypothesis testing (21). Full data are shown in **Dataset S4**. (B) Co-immunoprecipitation showing that RL1 interacts with DDB1. HEK-293s were stably transduced with RL1-V5 construct or controls. Input represents 1% of the sample. Proteins were detected with antibodies against V5 and DDB1. (C) Co-immunoprecipitation showing that interaction of RL1 and DDB1 is dependent largely on residues conserved between RL1 and UL145 (right panel, conserved residues shown in blue; UL145 residues required for interaction with DDB1 in red squares (19); RL1 residues required for interaction with DDB1 in green squares). HEK-293s were stably transduced with the indicated C-terminally V5-tagged RL1 constructs. Input represents 1% of the sample. Proteins were detected with antibodies against V5 and DDB1. (D) Immunoblot showing that SLFN11 downregulation is dependent on CUL4A, CUL4B and the adaptor protein DDB1. HFFF-TERTs stably expressing RL1-V5 or control were transfected for 48 h with siRNAs targeted against CUL4A, CUL4B, CUL4A/B, DDB1 or control. (E) Inhibition of CRL activity rescues SLFN11 levels. HFFF-TERTs stably transduced with RL1-V5 or control were treated with 1 μM MLN4924 for 24 h prior to harvest.

**Figure S4.**
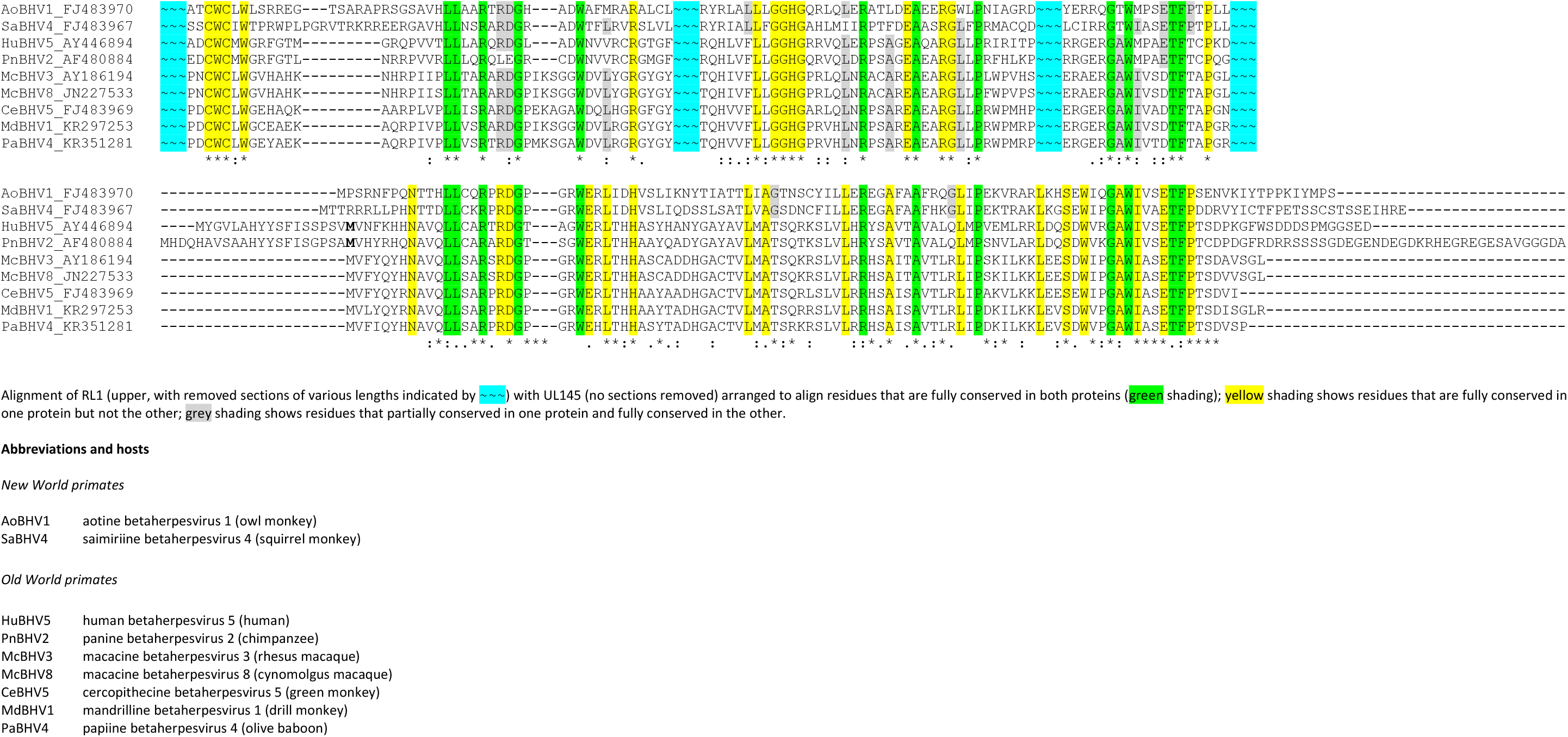
Alignment of RL1 with UL145.

To determine whether the CRL4 complex is required for RL1-mediated degradation of SLFN11, components of the complex were knocked down in HFFF-TERTs stably expressing RL1 or control. DDB1 knockdown completely restored SLFN11 (**Figures 3D and S5A**). SLFN11 was also rescued from degradation in the presence of MLN4924, which prevents the conjugation of NEDD8 on cullins (20), substantiating the requirement for the CRL4 complex in RL1-mediated SLFN11 degradation (**Figure 3E**). This suggests that RL1 may redirect the Cullin 4 ligase complex to degrade SLFN11, by acting as a viral DDB1-Cullin Accessory Factor (DCAF).

**Figure S5.**
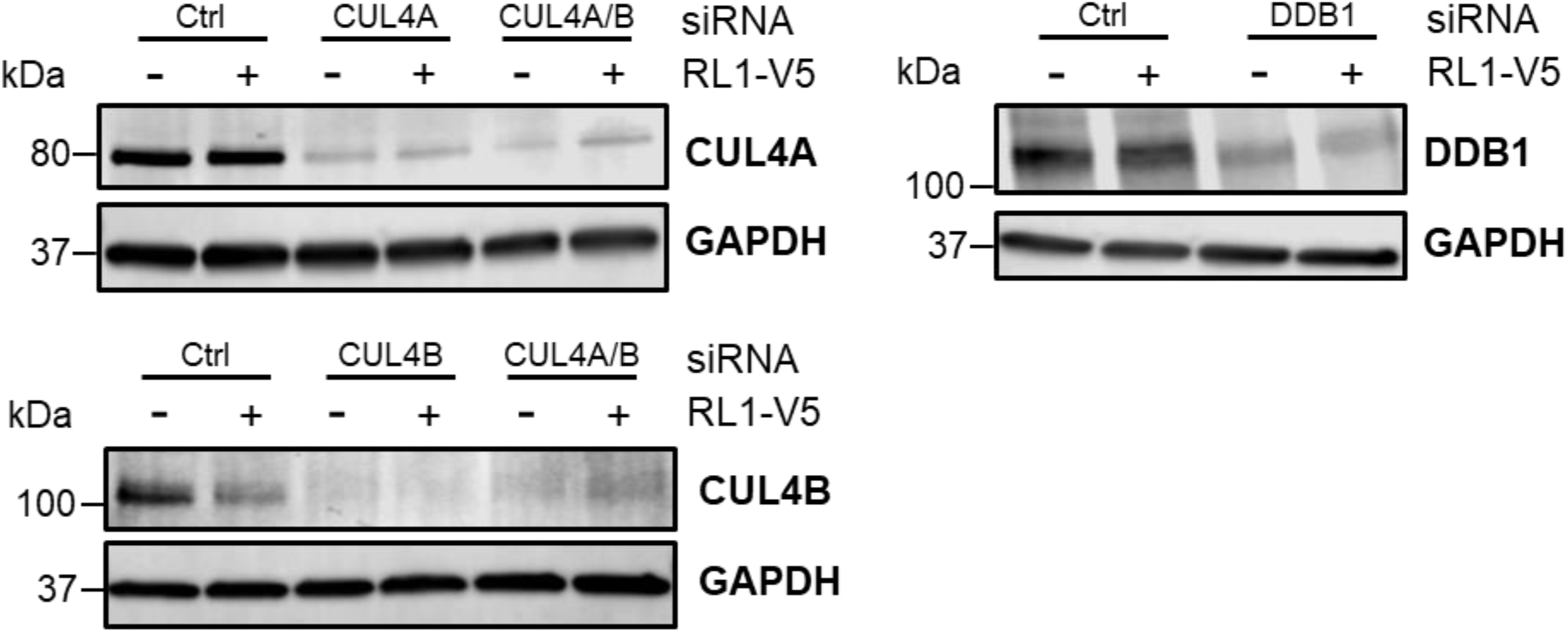
Demonstration of siRNA-mediated knockdown of CUL4A, CUL4B and DDB1, for the experiment shown in **Figure 3D**.

### SLFN11 restricts HCMV infection

We sought to determine whether SLFN11 restricts HCMV infection. SLFN11 depletion consistently and significantly increased HCMV replication in 4/4 independent HFFF-TERT cell lines stably knocked down for SLFN11, in terms of both number and size of plaques (**Figures 4A-B**). A decrease in the number of plaques was observed upon SLFN11 overexpression (**Figure 4C**). SLFN11 therefore represents a novel ARF for HCMV that acts to restrict significantly the spread of HCMV infection.

**Figure 4.**
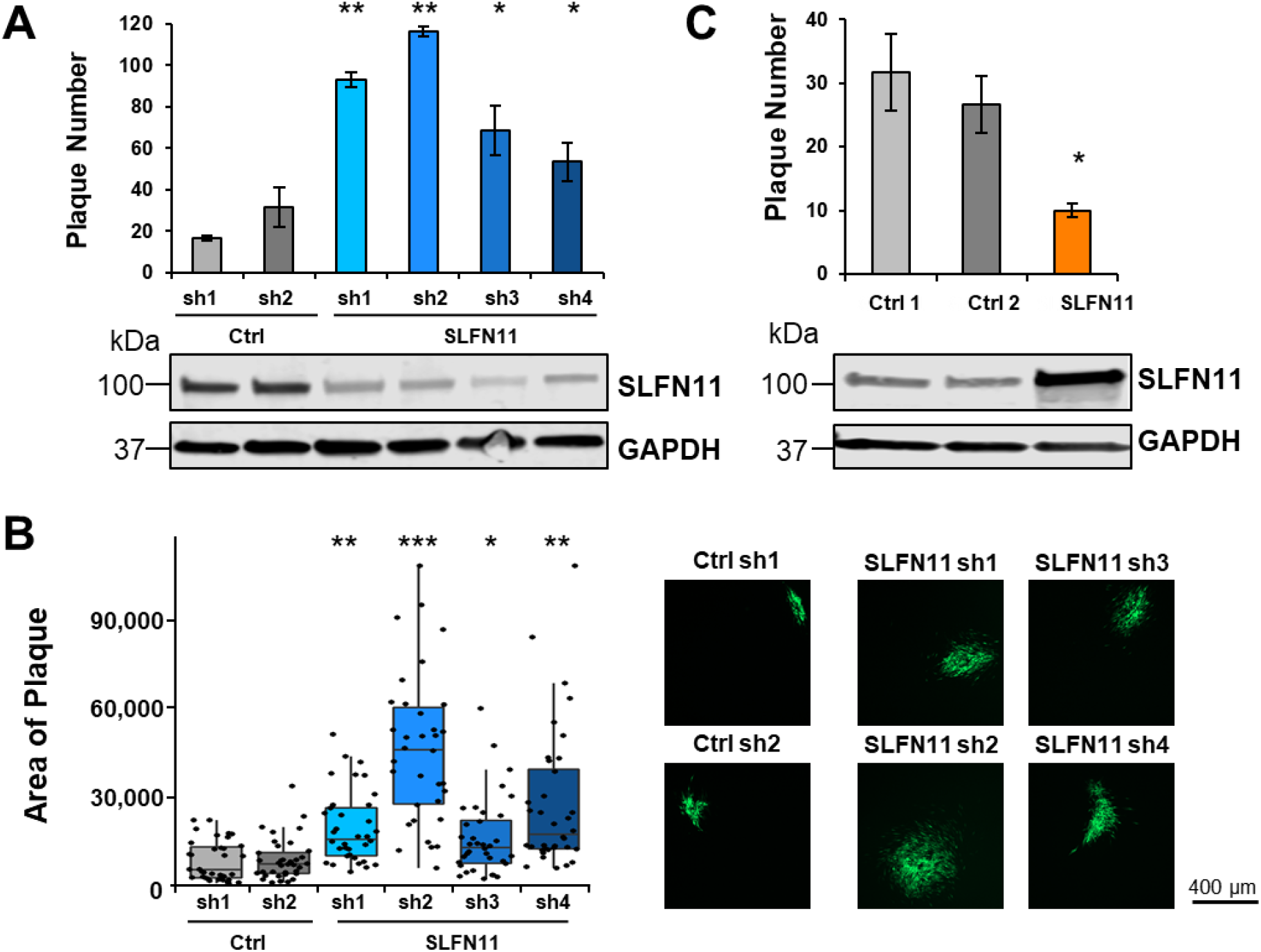
SLFN11 restricts HCMV infection. (A) SLFN11 restricts HCMV infection. HFFF-TERTs were stably transduced with shRNAs targeted against SLFN11 or control, and then infected in triplicate with AD169-GFP at MOI 0.005 under Avicel for 2 weeks before counting the number of plaques. A representative example of two experiments is shown, with error bars showing standard deviation from the mean. p-values were estimated using a two-tailed t-test (n=3). * p<0.05, ** p<0.0005. Immunoblot confirmed knockdown of SLFN11 (lower panel). (B) SLFN11 restricts cell-cell spread of HCMV. Plaque area was calculated using Fiji software (22) using pictures of plaques from the experiment described in (A). Representative examples are shown in the right panels. p-values were estimated using a non-parametric Mann-Whitney U test (n=30). * p < 0.0005, ** p<0.000005, *** p<5×10^−10^. (C) Confirmation that SLFN11 restricts HCMV infection. The experiment was conducted as described in (A), using HFFF-TERTs stably overexpressing SLFN11 or two independent control cell lines.

## Discussion

HCMV and other herpesviruses comprehensively modulate adaptive and innate immunity to facilitate their persistence, employing multiple viral proteins to target cellular factors for degradation (7). Although some viral proteins are expressed throughout the course of infection, others are temporally controlled and target a given host factor at a specific phase of viral replication (7, 11). The present study provides a systematic, searchable database that examines host protein regulation from the point of replication of the viral genome onwards, in addition to identifying which viral gene block targets each of >250 host factors.

The key roles of ARFs in protecting cell populations against HCMV are highlighted by the diversity of proteins with antiviral activity, with different factors affecting distinct steps of the HCMV replication cycle (reviewed in Schilling et al. (23)). Since description of protein components of promyelocytic leukemia bodies (PML, Sp100, hDaxx) as anti-HCMV ARFs, at least 15 additional ARFs have been identified, including HLTF, Zinc finger Antiviral Protein (ZAP), the cytidine deaminase APOBEC3A and the dNTP triphosphohydrolase SAMHD1. Some of these proteins exhibit antiviral activity against diverse viruses, whereas others, such as HLTF, have so far only been associated with restriction of HCMV.

We have now identified SLFN11 as a novel HCMV restriction factor, although the mechanism of restriction is yet to be determined. SLFN11 inhibits replication of lentiviruses in a codon usage-dependent manner, via its activity as a type II tRNA endonuclease (15, 16, 24, 25). Overall, HCMV genomes exhibit low codon usage bias, although the bias of individual coding sequences varies widely (26). Hu et al. (26) previously determined HCMV codon usage bias on a gene-by-gene basis. However, an analysis of their data using our temporal classification of HCMV protein expression (10) suggested that there is no systematic temporal codon usage bias of HCMV genes. It is possible that certain late-expressed viral genes that are poorly codon-optimised would be poorly translated in the absence of late RL1-mediated SLFN11 degradation. However, expression of poorly codon-optimised early expressed viral genes would presumably still be reduced. SLFN11 also inhibits translation of certain poorly codon-optimised human genes, in particular those specifying the serine/threonine kinases ATM and ATR (27). Both play key roles in the DNA damage response. HCMV requires ATM signaling for efficient replication, although the role of ATR signaling is presently unclear (reviewed in (28)). RL1 might thus prevent SLFN11-mediated repression of ATM/ATR to benefit viral replication. Further alternative mechanisms are suggested by the recent identification of SLFN5 as an ARF for herpes simplex virus 1 (HSV-1) and SLFN14 as an ARF for influenza virus. SLFN5 interacts with HSV-1 viral DNA to repress HSV-1 transcription (29), whereas SLFN14 promotes a delay in viral nucleoprotein translocation from cytoplasm to nucleus and enhances RIG-I mediated IFN-β signaling (30). These observations suggest that other components of the six-member human Schlafen family may act as restriction factors for HCMV, and that Schlafen proteins may more widely restrict other DNA and RNA viruses. Indeed, we found that SLFN5 was downregulated early during HCMV infection (Dataset S3), raising the intriguing possibility that the virus differentially regulates members of this important family to maximise viral replication.

Several viruses are now recognised to encode factors that degrade host protein targets by subverting cullins or their adaptor proteins, including hepatitis B, HIV, parainfluenza virus, bovine herpesvirus, murine gammaherpesvirus and CMVs (reviewed in (31, 32)). Including RL1, four CMV proteins have now been recognized to function in this manner, all via recruitment of CRL4 components: murine CMV-encoded M27 and HCMV-encoded UL35 and UL145 (7, 33–35). However, in our recent comprehensive HCMV interactome analysis (9), we detected six additional HCMV proteins that interact with CUL4A or CUL4B (RL12, US7, US34A, UL19, UL122 and UL135), two additional proteins interacting with DDB1 (UL19 and UL27), and three viral proteins interacting with other cullins (US30, UL26 and UL36). These data suggest that there are likely to be additional as yet uncharacterized mechanisms for HCMV -mediated cullin subversion, which may lead to degradation of additional host targets.

The presence of orthologs of RL1 and UL145 in the same positions and orientations in Old and New World monkey and ape cytomegalovirus genomes indicates that this pair of genes has existed for at least 40 million years. Furthermore, the conservation of amino acid residues required for DDB1 interaction suggest that the functions they serve are both ancient and essential for viral replication. Presumably, one or other of these genes developed first (perhaps by a now undetectable gene capture) and then duplicated. Sequences from early primate branches would be required to investigate the evolutionary history further, but these are presently lacking.

Our identification of RL1-mediated SLFN11 degradation provides the first evidence for direct viral antagonism of this important restriction factor, and might help to explain the evolution of SLFN11 under recurrent positive selection throughout primate development (25). Other mechanisms may also underlie this selection. Schlafen genes acquired by orthopoxviruses might inhibit their host counterparts, possibly by preventing cellular Schlafen multimerisation (25, 36). Certain flaviviruses might also encode anti-SLFN11 mechanisms, which could explain the differential susceptibility of West Nile, Zika and dengue viruses to SLFN11 effects (37). Additionally, sperm-egg interactions and meiotic drive can both result in strong signatures of recurrent positive selection, and some mammalian Schlafen genes have been implicated in sperm-egg incompatability (25, 36).

Only three drugs are commonly used in HCMV treatment, all exhibiting significant adverse effects and the risk of drug resistance. A novel therapeutic approach would be to prevent interaction of virally encoded immune antagonists with their cellular partners. The interaction of RL1 with SLFN11 is one example that could be inhibited for therapeutic effect. Other interactions involving distinct antiviral pathways could be targeted simultaneously to inhibit viral replication potently, for example between HCMV UL145 and HLTF. Alternatively, compounds that inhibit CRL function could be used in anti-HCMV therapy. It has been demonstrated that MLN4924 inhibits HCMV genome replication *in vitro* at nanomolar concentrations (31), but, to our knowledge, this compound has yet to be tested against HCMV in any clinical setting. Finally, our data are likely to identify further proteins that have roles in restricting infection by HCMV or other viruses.

## Materials and Methods

Extended materials and methods can be found in the supplementary information (SI)

### Viral infections for proteomic screens

HCMV strain Merlin was used in the PFA screen (38). Where indicated, cells were incubated with 300 μg/ml PFA (carrier: water) from the time of infection. For the block deletion mutant screen, 10 of the 11 block HCMV deletion mutants have been described previously (12). The ΔRL1-6 block deletion mutant was generated in the same fashion on the strain Merlin background lacking UL16 and UL18 and expressed a UL32-GFP reporter (wt2) (all viral recombinants used are shown in **Dataset S5A**). Detailed methods for whole cell lysate protein preparation and digestion, peptide labelling with TMT, HpRP fractionation, liquid chromatography-mass spectrometry, and data analysis are provided in the SI.

### Immunoprecipitation

Cells were harvested in lysis buffer, tumbled on a rotator and then clarified by centrifugation and filtration. After incubation with immobilised mouse monoclonal anti-V5 agarose resin, samples were washed and then subjected either to immunoblotting or to mass spectrometry (see SI).

### Plasmid construction and transduction

Lentiviral expression vectors encoding SLFN11, SLFN11-HA, or the V5-tagged viral proteins RL1, RL5A, RL6 and UL34 (control) were synthesised by PCR amplification and then cloned into Gateway vectors (50). V5-tagged RL1 point mutants were generated by PCR site-directed mutagenesis. For shRNA, two partially complementary oligonucleotides were annealed, and the resulting product was ligated into the pHR-SIREN vector. The primers and templates used are described in **Dataset S5C**. Stable cell lines were generated by transduction with lentiviruses produced via the transfection of HEK293T cells with the lentiviral expression vectors and helper plasmids.

### siRNA knockdown

HFFF-TERTs constitutively expressing RL1-V5 or control were transfected with pools of siRNAs for CUL4A, CUL4B, a mixture of CUL4A and CUL4B, DDB1 or non-targeting siRNAs (Dharmafect) with RNAiMAX (Thermo). Cellular lysates were harvested 48 h post transfection for immunoblotting.

### Immunoblotting

Protein concentration was measured in lysed cells using a bicinchoninic acid (BCA) assay. Aliquots (50 μg) of denatured, reduced protein was separated by SDS polyacrylamide gel electrophoresis (PAGE), transferred to a polyvinylidene difluoride (PVDF) membrane, and probed using the primary and secondary antibodies detailed in SI. Fluorescent signals were detected using the Odyssey CLx Imaging System (LI-COR), and images were processed and quantified using Image Studio Lite V5.2 (LI-COR).

### Plaque assay

HFFF-TERTs stably expressing shRNA constructs targeted against SLFN11 or control, or overexpressing SLFN11 or control, were infected in triplicate at MOI 0.005 with RCMV-288 (strain AD169 expressing enhanced green fluorescent protein under the control of the HCMV β-2.7 early promoter) (39). The medium was then replaced with a 1:1 (v/v) mixture of 2 x DMEM and Avicel (2% (w/v) in water). This mixture was removed 2 weeks after infection and the cells were washed then fixed in 4% (w/v) paraformaldehyde. The number of plaques per well was counted on the basis of GFP fluorescence. Plaque area was calculated using Image J Fiji software.

## Acknowledgments

We are grateful to Prof. Steve Gygi for providing access to the “MassPike” software pipeline for quantitative proteomics. This work was supported by a Wellcome Trust Senior Clinical Research Fellowship (108070/Z/15/Z) to MPW, MRC Project Grants (MR/P001602/1) to ECYW, (MR/S00971X/1, MR/V000489/1) to RJS and ECYW, and an MRC Programme Grant (MC_UU_12014/3) to AJD. This study was additionally supported by the Cambridge Biomedical Research Centre, UK.

## Supplementary information

### Materials and Methods

#### Cells and cell culture

Primary human fetal foreskin fibroblast cells (HFFFs), HFFFs immortalised with human telomerase (HFFF-TERTs), HEK-293T and HEK-293 cells were grown in Dulbecco’s modified Eagle’s medium (DMEM) supplemented with 10% (v/v) foetal bovine serum (FBS) and 100 IU/ml penicillin / 0.1 mg/ml streptomycin (DMEM+FBS) at 37°C in 5% CO_2_. HFFFs and HFFF-TERTs have been tested at regular intervals since isolation to confirm that human leukocyte antigen (HLA) and MHC Class I Polypeptide-Related Sequence A (MICA) genotypes, cell morphology and antibiotic resistance properties are consistent with the original cells. In addition, growth of the HCMV strain used is limited to human fibroblasts (dermal or foreskin), further limiting the chances that the cells have been contaminated with another cell type.

For SILAC immunoprecipitations, cells were grown for seven divisions in DMEM supplemented with 10% (v/v) dialysed FBS, 100 IU/ml penicillin / 0.1 mg/ml streptomycin, 280 mg/l L-proline and either light (Arg 0, Lys 0), medium (Arg 6, Lys 4) or heavy (Arg 10, Lys 8) amino acids at 50 mg/l. Where indicated, cells were treated with 10 mM MG132 (Merck) for 12 h, or 1 μM MLN4924 (Abcam, ab216470) for 24 h prior to harvesting.

#### Viruses

The parental virus (RCMV1111) used was derived by transfection of a bacterial artificial chromosome (BAC) clone of HCMV strain Merlin, the genome of which is designated the reference HCMV sequence by the National Center for Biotechnology Information and was sequenced after three passages in vitro (1, 2). RCMV1111 contains point mutations in two genes (RL13 and UL128) that enhance replication in fibroblasts (2). Ten of the 11 HCMV block deletion mutants used have been described previously (3) and were generated on a strain Merlin background (wt1) or wt1 that lacked UL16 and UL18 and expressed a UL32-GFP reporter (wt2). The ΔRL1-6 block deletion mutant was generated in the same fashion on the wt2 background and validated by whole genome sequencing. RCMV288 was used for plaque assays, and is based on HCMV strain AD169 with one copy of the EGFP (enhanced green fluorescent protein) gene inserted in one copy of the HCMV long repeat under the control of the HCMV RNA2.7 early promoter (between nucleotides 4576 to 2154 with respect to the strain AD169 genomic sequence) (4). All viral recombinants used are described in **Dataset S5A**.

#### Virus infections

For the Phosphonoformate (PFA) screen, 1.5 x 10^7^ HFFFs were plated in a 150 cm^2^ flask 24 h prior to infection with HCMV strain Merlin at MOI 10. Where indicated, cells were incubated with 300 μg/ml PFA from the time of infection. For the block deletion mutant screen, 1 x 10^6^ HFFF-TERTs were plated in a 25 cm^2^ flask 24 h prior to infection with HCMV strain Merlin at MOI 5 as previously described (3). Mock infections were performed identically but with DMEM instead of viral stock. The zero time point (0 h) was considered to be the time at which cells first came into contact with virus. Cells were incubated with virus for 2 h at 37 °C on a rocking platform, and then the medium was replaced with DMEM+FBS.

#### Whole cell lysate protein digestion

Cells were washed twice with phosphate buffered saline (PBS), and 250 μl lysis buffer (6 M guanidine, 50 mM HEPES pH 8.5) was added. Cell lifters (Corning) were used to scrape the cells into the lysis buffer, and the suspension was transferred into a 1.5 ml Eppendorf tube, vortexed extensively, and sonicated. Cell debris was removed by centrifuging twice at 21,000 g for 10 min. Half of each sample was kept for subsequent analysis by immunoblot where required. For the other half, dithiothreitol (DTT) was added to a final concentration of 5 mM and samples were incubated for 20 min at room temperature. Cysteine residues were alkylated with 14 mM iodoacetamide and incubated at room temperature for 20 min in the dark. Excess iodoacetamide was quenched with 5mM DTT for 15 mins. Samples were diluted with 200 mM HEPES pH 8.5 to bring the guanidine concentration to1.5 M and digested at room temperature for 3 h with LysC protease at a protease-to-protein ratio of 1:100. Samples were further diluted with 200 mM HEPES pH 8.5 to bring the guanidine concentration to 0.5 M. Trypsin was then added at a protease-to-protein ratio of 1:100, and the reactions were incubated overnight at 37°C. Reactions were quenched with 5% (v/v) formic acid and centrifuged at 21,000 g for 10 min to remove undigested protein. Peptides were subjected to C18 solid-phase extraction (SPE, Sep-Pak, Waters) and vacuum-centrifuged to near-dryness.

#### Peptide labelling with tandem mass tags (TMTs)

In preparation for TMT labeling, desalted peptides were dissolved in 200 mM HEPES pH 8.5. Peptide concentration was measured by microBCA (Pierce). TMT reagents (0.8 mg) were dissolved in 43 μl anhydrous acetonitrile and 3 μl added to 25 μg of peptide at a final acetonitrile concentration of 30% (v/v). Sample labelling was as indicated in **Dataset S5B**. Following incubation at room temperature for 1 h, the reaction was quenched with hydroxylamine to a final concentration of 0.3% (v/v). The nine TMT-labelled samples that constituted the PFA screen were combined at a ratio of 1:1:1:1:1:1:1:1:1, and the ten samples that constituted the block deletion mutant screen were combined at a ratio of 1:1:1:1:1:1:1:1:1:1. Combined samples were vacuum-centrifuged to near dryness and subjected to C18 SPE (Sep-Pak, Waters). An unfractionated ‘singleshot’ sample was analysed initially to ensure similar peptide loading across each TMT channel and avoid the need for excessive electronic normalization. As all normalisation factors were >0.5 and <2, data for the PFA singleshot experiment was analysed with data for the corresponding fractions to increase the overall number of peptides quantified.

#### Offline high pH reversed phase (HpRp) fractionation and LC-MS/MS/MS for the PFA experiment

Combined, TMT-labeled peptide samples were fractionated using an Agilent 300Extend C18 column (5 μm particles, 4.6 mm ID, 220 mm length) and an Agilent 1100 quaternary pump equipped with a degasser and a photodiode array detector (220 and 280 nm, ThermoFisher, Waltham, MA). Peptides were separated with a linear gradient of 5-35% (v/v) acetonitrile in 10 mM ammonium bicarbonate pH 8 over 60 min. Fractions were recombined orthogonally in a checkerboard fashion, combining alternate wells from each column of the plate into a single fraction, and commencing combination of adjacent fractions in alternating rows. Wells were excluded prior to the start or after the cessation of elution of peptide-rich fractions, as identified from the UV trace. This yielded two sets of 12 combined fractions, A and B. All 12 set A fractions were dried in a vacuum centrifuge, desalted using a StageTip (5) and resuspended in 10 μl MS solvent (4% (v/v) MeCN, 5% (v/v) formic acid) prior to LC-MS/MS/MS.

For LC/MS/MS/MS, peptides were separated on a 75 μm inner diameter microcapillary column packed with 0.5 cm of Magic C4 resin (5 μm, 100 Å, Michrom Bioresources) followed by approximately 20 cm of GP118 resin (1.8 μm, 120 Å, Sepax Technologies). Peptides were separated using a 3 h linear gradient (WCL samples) or 2 h gradient (PM samples) of 6-30% (v/v) acetonitrile in 0.125% (v/v) formic acid at a flow rate of 300 nl/min. Each analysis used an MS3-based TMT method (6, 7). The scan sequence began with an MS1 spectrum (Orbitrap analysis, resolution 120,000, 400-1400 Th, AGC target 2 × 10^5^, maximum injection time 200 ms). ‘Top speed’ (2s) was selected for MS2 analysis, which consisted of CID (quadrupole ion trap analysis, AGC 4 × 10^3^, NCE 35, maximum injection time 150 ms). The top ten precursors were selected for MS3 analysis, in which precursors were fragmented by HCD prior to Orbitrap analysis (NCE 55, max AGC 5 × 10^4^, maximum injection time 250 ms, isolation specificity 0.5 Th, resolution 60,000) (6).

#### Offline HpRp fractionation and LC-MS/MS/MS for the block deletion mutant experiment

TMT-labelled tryptic peptides were subjected to HpRP fractionation using an Ultimate 3000 RSLC UHPLC system (Thermo Fisher Scientific) equipped with a 2.1 mm internal diameter (ID) x 25 cm long, 1.7 mm particle Kinetix Evo C18 column (Phenomenex). The mobile phase consisted of A: 3% (v/v) acetonitrile (MeCN), B: 100% MeCN and C: 200 mM ammonium formate pH 10. Isocratic conditions were 90% A/10% C, and C was maintained at 10% throughout the gradient elution. Separations were conducted at 45°C. Samples were loaded at 200 ml/min for 5 min. The flow rate was then increased to 400 ml/min over 5 min, after which the gradient elution proceed as follows: 0-19% B over 10 min, 19-34% B over 14.25 min, 34-50% B over 8.75 min, finally washing with 90% B over 10 min. UV absorbance was monitored at 280 nm, and 15 s fractions were collected into 96 well microplates using the integrated fraction collector. Fractions were recombined orthogonally in a checkerboard fashion (as described above), combining alternate wells from each column of the plate into a single fraction, and commencing combination of adjacent fractions in alternating rows. Wells were excluded prior to the start or after the cessation of elution of peptide-rich fractions, as identified from the UV trace. This yielded two sets of 12 combined fractions, A and B. All 24 fractions were dried in a vacuum centrifuge and resuspended in 10 μl MS solvent (4% MeCN, 5% formic acid) prior to LC-MS/MS/MS.

For LC/MS/MS/MS, the loading solvent was 0.1% (v/v) TFA and the analytical solvents were A: 0.1% (v/v) formic acid (FA) and B: MeCN + 0.1% (v/v) FA. All separations were carried out at 55°C. Samples were loaded at 10 ml/min for 5 min in loading solvent before beginning the analytical gradient. The following gradient was used, whereby A was used to dilute B at the percentage shown: 3-5.6% B over 4 min, 5.6-32% B over 162 min, followed by a 5 min wash with 80% B, a 5 min wash with 90% B and equilibration at 3% B for 5 min. Each analysis used a MultiNotch MS3-based TMT method (McAlister et al., 2014). The following settings were used: MS1: 400-1400 Th, Quadrupole isolation, 120,000 Resolution, 2×10^5^ AGC target, 50 ms maximum injection time, ions injected for all parallisable time. MS2: Quadrupole isolation at an isolation width of m/z 0.7, CID fragmentation (NCE 30) with ion trap scanning out in rapid mode from m/z 120, 1×10^4^ AGC target, 70 ms maximum injection time, ions accumulated for all parallisable time in centroid mode. MS3: in Synchronous Precursor Selection mode the top 10 MS2 ions were selected for HCD fragmentation (NCE 65) and scanned in the Orbitrap at 50,000 resolution with an AGC target of 5×10^4^ and a maximum accumulation time of 150 ms, and ions were not accumulated for all parallelisable time. The entire MS/MS/MS cycle had a target time of 3 s. Dynamic exclusion was set to ±10 ppm for 90 s. MS2 fragmentation was trigged on precursors 5×103 counts and above.

#### Immunoprecipitation, protein digestion and LC/MS/MS for interaction analysis

Cells were harvested in lysis buffer (50 mM Tris pH 7.5, 300 mM NaCl, 0.5% (v/v) NP40, 1 mM DTT and Roche protease inhibitor cocktail), tumbled for 15 min at 4°C and then centrifuged at 16,100 g for 20 min at 4°C. Lysates were clarified by filtration through a 0.7 μm filter and incubated for 3 h with immobilised mouse monoclonal anti-V5 agarose resin. Samples were washed seven times with lysis buffer and then seven times with PBS pH 7.4. Proteins bound to the anti-V5 resin were eluted twice by adding 200 ml of 250 mg/ml V5 peptide (Alpha Diagnostic International) in PBS at 37°C for 30 min with agitation. Finally, proteins were precipitated with 20% (v/v) TCA, washed once with 10% (v/v) TCA, washed three times with cold acetone and dried to completion using a centrifugal evaporator. Samples were resuspended in digestion buffer (50 mM Tris pH 8.5, 10% (v/v) MeCN, 1 mM DTT, 10 ug/ml trypsin) and incubated overnight at 37°C with agitation. The reaction was quenched with 50% (v/v) FA, subjected to C18 solidphase extraction, vacuum-centrifuged to complete dryness, and resuspended in 10 μl MS solvent (4% (v/v) MeCN, 5% (v/v) formic acid) prior to LC-MS/MS. For co-immunoprecipitation, the above protocol was followed except that the number of washes was reduced to three with lysis buffer and two with PBS. Proteins bound to the anti-V5 resin were eluted by adding 40 μl of 250 μg/ml V5 peptide in PBS at 37°C for 30 min with agitation. Eluates were separated by SDS-PAGE as described below.

For LC/MS/MS, the loading solvent was 3% MeCN, 0.1% FA and the analytical solvents were A: 0.1% (v/v) FA and B: MeCN + 0.1% (v/v) FA. All separations were carried out at 55°C. Samples were loaded at 5 ml/min for 5 min in loading solvent before beginning the analytical gradient. The following gradient was used: 3-40% B over 29 min followed by a 3 min wash in 95% B and equilibration in 3% B for 10 min. The following settings were used: MS1: 300-1500 Th, 120,000 resolution, 4×10^5^ AGC target, 50 ms maximum injection time. MS2: Quadrupole isolation at an isolation width of m/z 1.6, HCD fragmentation (NCE 35) with fragment ions scanning in the Orbitrap from m/z 110, 5×10^4^ AGC target, 60 ms maximum injection time, ions accumulated for all parallelisable time. Dynamic exclusion was set to ±10 ppm for 60 s. MS2 fragmentation was trigged on precursors 5×10^4^ counts and above.

#### Plasmid construction

V5-tagged RL1, RL5A, RL6 and UL34 (as a control) were amplified by PCR from adenoviral templates containing the tagged viral gene under the control of the HCMV major immediate early promoter (MIEP). Primers were designed to recognise the 3’ end of the MIEP (forward) and the V5 tag (reverse) (**Dataset S5C**). SLFN11 was amplified from HFFF-TERT cDNA using primers designed to recognise the 5’ and 3’ ends of the gene. For the HA-tagged construct, the reverse primer contained a 6 bp linker followed by the coding sequence for an HA tag and a stop codon. A control construct was prepared by annealing two oligonucleotides detailed in **Dataset S5C**. Both primers and oligonucleotides had flanking Gateway attB sequences for the purposes of cloning (**Dataset S5C**). PCR employed PfuUltra II Fusion HS DNA polymerase (Agilent). Constructs were subsequently cloned into lentiviral destination vector pHAGE-pSFFV using the Gateway system (Thermo Scientific).

To generate shRNA constructs, two partially complementary oligonucleotides were annealed, with all sequences shown in **Dataset S5C**. The resulting product was ligated as a BamHI–EcoRI fragment into the pHR-SIREN vector (a gift from Prof. Paul Lehner, University of Cambridge) using T4 ligase (Thermo Scientific). All constructed plasmids were transformed into 5-alpha Competent *E. coli* (NEB) and selected on antibiotic-containing luria broth (LB) agar plates. All plasmid inserts were sequenced fully to check for mutations. Two different non-targeting control shRNA sequences are shown in **Dataset S5C**, which also lists the sequences of all primers and oligonucleotides.

#### Site-directed mutagenesis

A method based on PCR overlap extension was used to generate point mutations in the coding sequence of RL1. Primer sequences spanning the target region were generated that incorporated the desired sequence changes in both forward and reverse orientations. These, along with primers that would anneal at the 5’ and 3’ ends of the full-length RL1 coding sequence (RL1F and RL1R, respectively) were used to amplify two fragments of RL1, each incorporating the point mutation (**Dataset S5C**). Fragments were purified and assembled into a full-length mutant RL1 coding sequence by a second round of PCR using only RL1F and RL1R. The product was then purified and subcloned as described above.

#### Stable cell line production

Lentiviral particles were generated by transfection of HEK-293T cells with the lentiviral transfer vector plus four helper plasmids (VSVG, TAT1B, MGPM2, CMV-Rev1B), using TransIT-293 transfection reagent (Mirus) according to the manufacturer’s recommendations. Viral supernatant was typically harvested 48 h after transfection, cell debris was removed with a 0.22 μm filter, and target cells were transduced for 48 h and then subjected to antibiotic selection for two weeks.

#### siRNA knockdown

24 h prior to transfection, 2.5 x 10^5^ HFF-TERTs constitutively expressing RL1-V5 or control were plated in 6-well plates. Cells were transfected with a pool of CUL4A siRNAs (L-012610-00, Dharmafect), a pool of CUL4B siRNAs (L-017965-00, Dharmafect), a pool of DDB1 siRNAs (L-012890-00, Dharmafect), a 1:1 mixture of CUL4A and CUL4B pools, or a pool of non-targeting siRNAs (D-001810-10, Dharmafect) with RNAiMAX (13778030, Thermo), giving a final siRNA concentration of 50 nM. Cellular lysates were harvested for immunoblotting at 48 h post transfection.

#### Immunoblotting

Cells were lysed with RIPA buffer (Cell Signaling) containing complete protease inhibitor cocktail (Roche), and protein concentrations were measured using a bicinchoninic acid assay (BCA, Pierce). Lysates were reduced with 6 x protein loading dye (375 mM Tris pH 6.8, 12% (w/v) sodium dodecyl sulfate (SDS), 30% (v/v) glycerol, 0.6 M DTT, 0.06% bromophenol blue) for 5 min at 95°C. An amount (50 μg) of protein for each sample was separated by polyacrylamide gel electrophoresis (PAGE) using 4-15% (w/v) TGX precast protein gels (Bio-rad) and then transferred to polyvinylidene fluorid (PVDF) membranes using a trans-blot system (Bio-rad). The following primary antibodies were used: anti-glyceraldehyde 3-phosphate dehydrogenase (GAPDH) (1:10,000, #MAB5718, R&D Systems), anti-V5 (1:2000, #MA5-15253, Thermo), anti-V5 (1:1000, #D3H8Q, CST), anti-SLFN11 (1:500, #HPA023030, Atlas Antibodies), anti-CUL4A (1:1000, #2699S, CST), anti-DDB1 (1:1000, #5428S, CST) and anti-CUL4B (1:1000, #ab157103, Abcam). Secondary antibodies were IRDye 680RD goat anti-mouse (925-68070, LI-COR), IRDye 680RD goat anti-rabbit (926-68071, LI-COR), IRDye 800CW goat anti-mouse (926-32210, LI-COR) and IRDye 800CW goat anti-rabbit (925-32211, LI-COR).

#### Plaque assay

1.3 x 10^5^ HFFF-TERTs stably expressing shRNA constructs targeted against SLFN11 or control, or overexpressing SLFN11 or control were plated in 12-well plates in triplicate 24 h prior to infection with AD169-GFP at MOI 0.005. Cells were incubated with virus for 2 h prior to replacing the medium with a 1:1 (v/v) mixture of 2 x DMEM and Avicel (2% w/v in water, FMC BioPolymer). At 2 weeks post infection, the DMEM/Avicel mixture was removed and the cells were washed three times with PBS before fixation in 4% (w/v) paraformaldehyde. The number of plaques per well was counted on the basis of GFP fluorescence and the average plaque number was calculated across the three wells for each condition.

#### Plaque size analysis

Images of 12 plaques per well were taken using a Zeiss AxioObserver inverted fluorescence microscope at 5 x magnification. The first 12 plaques encountered that were suitable for imaging were used. Suitable plaques were those that fitted entirely in the field of view, were not located against the edge of the well, and were the only plaque within the field of view. Images were converted into greyscale and plaque area was calculated using Image J Fiji.

#### Data analysis

Mass spectra were processed using a Sequest-based software pipeline for quantitative proteomics, “MassPike”, through a collaborative arrangement with Professor Steven Gygi’s laboratory at Harvard Medical School. MS spectra were converted to mzXML using an extractor built upon Thermo Fisher’s RAW File Reader library (version 4.0.26). In this extractor, the standard mzxml format has been augmented with additional custom fields that are specific to ion trap and Orbitrap mass spectrometry and essential for TMT quantitation. These additional fields include ion injection times for each scan, Fourier transform-derived baseline and noise values calculated for every Orbitrap scan, isolation widths for each scan type, scan event numbers, and elapsed scan times. This software is a component of the MassPike software platform and is licensed by Harvard Medical School.

A combined database was constructed from (a) the human Uniprot database (26 January 2017), (b) the HCMV strain Merlin Uniprot database, (c) all additional non-canonical human cytomegalovirus ORFs described by Stern-Ginossar et al (8), (d) a six-frame translation of HCMV strain Merlin filtered to include all potential ORFs of ≥8 amino acids (delimited by stop-stop rather than requiring ATG-stop) and (e) common contaminants such as porcine trypsin and endoproteinase LysC. ORFs from the six-frame translation (6FT-ORFs) were named as follows: 6FT_Frame_ORFnumber_length, where Frame is numbered 1-6, and length is the length in amino acids. The combined database was concatenated with a reverse database composed of all protein sequences in reversed order. Searches were performed using a 20 ppm precursor ion tolerance. Fragment ion tolerance was set to 1.0 Th. TMT tags on lysine residues and peptide N termini (229.162932 Da) and carbamidomethylation of cysteine residues (57.02146 Da) were set as static modifications, while oxidation of methionine residues (15.99492 Da) was set as a variable modification. For SILAC analysis, the following variable modifications were used: heavy lysine (8.01420 Da), heavy arginine (10.00827 Da), medium lysine (4.02511 Da), medium arginine (6.02013 Da). SILAC-only searches were performed in the same manner, omitting the TMT static modification.

To control the fraction of erroneous protein identifications, a target-decoy strategy was employed (9). Peptide spectral matches (PSMs) were filtered to an initial peptide-level false discovery rate (FDR) of 1% with subsequent filtering to attain a final protein-level FDR of 1%. PSM filtering was performed using a linear discriminant analysis, as described previously (9). This distinguishes correct from incorrect peptide identifications (IDs) in a manner analogous to the widely used Percolator algorithm (https://noble.gs.washington.edu/proj/percolator/), by employing a distinct machine learning algorithm. The following parameters were considered: XCorr, ΔCn, missed cleavages, peptide length, charge state, and precursor mass accuracy.

Protein assembly was guided by principles of parsimony to produce the smallest set of proteins necessary to account for all observed peptides (algorithm described in (9). Where all PSMs from a given HCMV protein could be explained either by a canonical gene or non-canonical ORF, the canonical gene was picked in preference.

In a few cases, PSMs assigned to a non-canonical gene or 6FT-ORF were a mixture of peptides from the canonical protein and the ORF. This most commonly occurred where the ORF was a 5’-terminal extension of the canonical gene (thus meaning that the ORF was the simplest explanation for the observed peptides). In these cases, the peptides corresponding to the canonical protein were separated from those unique to the ORF, generating two separate entries. In a single case, PSMs were assigned to the 6FT-ORF 6FT_6_ORF1202_676aa, which is a 5’-terminal extension of the non-canonical ORF ORFL147C. The principles described above were used to separate these two ORFs.

Proteins were quantified by summing TMT reporter ion counts across all matching peptide-spectral matches using “MassPike”, as described previously (McAlister et al., 2014). Briefly, a 0.003 Th window around the theoretical m/z of each reporter ion (126, 127n, 127c, 128n, 128c, 129n, 129c, 130n, 130c, 131n) was scanned for ions, and the maximum intensity nearest to the theoretical m/z was used. The primary determinant of quantitation quality is the number of TMT reporter ions detected in each MS3 spectrum, which is directly proportional to the signal-to-noise (S:N) ratio observed for each ion. Conservatively, every individual peptide used for quantitation was required to contribute sufficient TMT reporter ions (minimum of ~500 per spectrum) so that each on its own could be expected to provide a representative picture of relative protein abundance (10). An isolation specificity filter with a cutoff of 50% was additionally employed to minimise peptide co-isolation (10). Peptide-spectral matches with poor quality MS3 spectra (more than 9 TMT channels missing and/or a combined S:N ratio of <100 across all TMT reporter ions) or no MS3 spectra at all were excluded from quantitation. Peptides meeting the stated criteria for reliable quantitation were then summed by parent protein, in effect weighting the contributions of individual peptides to the total protein signal based on their individual TMT reporter ion yields. Protein quantitation values were exported for further analysis in Excel.

For protein quantitation, reverse and contaminant proteins were removed, then each reporter ion channel was summed across all quantified proteins and normalised assuming equal protein loading across all channels. For further analysis and display in Figures, fractional TMT signals were used (i.e. reporting the fraction of maximal signal observed for each protein in each TMT channel, rather than the absolute normalized signal intensity). This effectively corrected for differences in the numbers of peptides observed per protein. For all TMT or SILAC experiments, normalised S:N values are presented in supplementary tables, assuming equal protein loading across all samples. As it was not possible confidently to assign peptides to only two HLA-A, HLA-B or HLA-C alleles, S:N values were further summed to give a single combined result for HLA-A, HLA-B or HLA-C.

Three block viral gene-deletion screens were conducted as described above. For each protein in each screen, a mean (μ) and standard deviation (σ) of all normalised S:N values was calculated. In each case, the maximum (x) value was omitted. For example, for SLFN11 in Figure 2C, μ and σ were calculated using values for wt1, wt2, RL10-UL1, RL11-UL11, UL2-UL11, UL13-UL20, UL/b’, US1-US11, US12-US17, US18-US22, US27-US28, US29-US34A but not the maximum RL1-RL6. The formula z = (x – μ) / σ was then applied to calculate a z-score. Fold change (FC) compared to wild-type (wt) infection was calculated from normalised S:N values using FC = x/wt1. For each experiment, a given protein was initially assigned to the block corresponding to the TMT channel with the maximum S:N. To combine results to assign an overall block to each protein, if the protein was quantified in two or three screens and assigned to the same block, z-scores and fold changes were averaged. If the protein was only quantified in one of the three screens, the block assignment, z-score and fold change from that screen were used. Otherwise, it was not considered possible to assign an overall gene block for that protein.

Hierarchical centroid clustering based on uncentered Pearson correlation, and k-means clustering were performed using Cluster 3.0 (Stanford University) and visualised using Java Treeview (http://jtreeview.sourceforge.net). p-values for protein fold change were estimated using the method of Significance A, calculated in MaxQuant and corrected for multiple hypothesis testing using the method of Benjamini-Hochberg (11). Multiple sequence alignment was performed using Clustal Omega (http://www.ebi.ac.uk/Tools/msa/clustalo/) provided by EMBL-EBI. Cited values for nucleotide and amino acid sequence identity do not include alignment gaps.

#### Data and materials availability statement

The mass spectrometry raw files will be deposited in the ProteomeXchange Consortium via the PRIDE partner repository (12). All materials described in this manuscript, and any further details of protocols employed can be obtained on request from the corresponding author by email to.

#### Dataset S1 (separate xls. file)

A. Proteins identified by the PFA screen as downregulated >3-fold by 96 hpi and ‘rescued’ >2-fold by PFA, with both fold changes significant at p<0.1 (see Figure 1B).

B. Enrichment of functional pathways within proteins identified in (A), compared to all proteins quantified from this screen.

C. Proteins identified by the PFA screen as upregulated >3-fold by 96 hpi yet inhibited >2-fold by PFA, with both fold changes significant at p<0.1 (see Figure 1B).

#### Dataset S2 (separate xls.file).

Predicted block of viral genes targeting 254 human proteins.

#### Dataset S3 (separate xls. file)

Interactive spreadsheet of all data in the manuscript. The “Plotter” worksheet generates graphs for all the human and viral proteins quantified, and visualises statistics. The “Data” worksheet shows minimally annotated protein data, for which the only modifications are formatting, normalization, reassignment of non-canonical HCMV ORFs and combination of peptide data for HLA- A, -B and -C alleles into single combined results, as described in Materials and Methods. The “Lookup” worksheet is used to generate graphs shown in “Plotter”.

#### Dataset S4 (separate xls. file)

Full results of the SILAC immunoprecipitation of RL1, RL5A and RL6 (see also **Figure 3A**).

#### Dataset S5 (separate xls. file)

**A.** Details of viruses and viral block deletion recombinants used in this study.

**B.** Details of TMT labelling for each proteomic experiment.

**C.** Details of oligonucleotides for gene knock-down, overexpression or mutagenesis.

